# Underlying selection for the diversity of Spike protein sequences of SARS-CoV-2

**DOI:** 10.1101/2021.10.22.465411

**Authors:** Manisha Ghosh, Surajit Basak, Shanta Dutta

**Affiliations:** Division of Bioinformatics, ICMR-National Institute of Cholera and Enteric Diseases, Kolkata, India; Division of Bacteriology, ICMR-National Institute of Cholera and Enteric Diseases, Kolkata, India

**Keywords:** Amino acid usage, Hydrophobicity, Correspondence analysis, Molecular docking, Spike protein sequence

## Abstract

The global spread of SARS-CoV-2 is fast moving and has caused a worldwide public health crisis. In the present manuscript we analyzed spike protein sequences of SARS-CoV-2 genomes to assess the impact of mutational diversity. We observed from amino acid usage patterns that spike proteins are associated with a diversity of mutational changes and most important underlying cause of variation of amino acid usage is the changes in hydrophobicity of spike proteins. The changing patterns of hydrophobicity of spike proteins over time and its influence on the receptor binding affinity provides crucial information on the SARS-CoV-2 interaction with human receptor. Our results also show that spike proteins have evolved to prefer more hydrophobic residues over time. The present study provides a comprehensive analysis of molecular sequence data to consider that mutational variants might play a crucial role in modulating the virulence and spread of the virus and has immediate implications for therapeutic strategies.

## Introduction

Coronaviruses are members of the family Coronaviridae. They are single-stranded, positive-sense RNA viruses that cause widespread respiratory, gastrointestinal and neurological clinical symptoms ^1,2^. To survive under immunological pressure within humans the virus accumulates mutations to outwit the immune system ^3^. These mutations may lead to change the virulence of the virus and its infectivity ^4^. Therefore, it is important to analyze the mutational pattern in order to ascertain virus evolutionary dynamics.

Gene sequence data from pathogen genome has been widely recognized as an important tool to study the infection dynamics ^5,6^. Coronavirus replication is error prone as compared to other RNA viruses and the estimated mutation rate is 4×10^−4^ nucleotide substitutions/site/year ^7^. Wang et al. ^8^ reported 13 mutations in SARS-CoV-2 genome from the genome sequences submitted till February 2020. In another study, 93 mutations were identified across the SARS-CoV-2 genome which includes three mutations in RBD of S protein demanding further study to understand the impact of these mutations on antigenicity of the SARS-CoV-2 ^9^. van Dorp et al. ^10^ studied 7666 public genome assemblies of SARS-CoV-2 and identified invariant and diversified regions of the genome. They observed 198 recurrent mutations across the genome when compared with reference genomes of Wuhan-Hu-1 (accession IDs C_045512.2 and EPI_ISL_402125). Earlier reports found that most (80%) of the mutations were non-synonymous at the protein level ^9,11^. More recent studies suggest that the D614G variant is close to reaching fixation around the world ^12^. Groves et al. (2021) suggested that mutations in spike proteins which are associated with higher viral loads may lead to a more open conformation enhancing the binding of the virus spike to the ACE2 receptor ^13^.

Coronavirus uses spike proteins for primary interaction with human host. Spike protein binds with the human cellular receptor angiotensin converting enzyme 2 (ACE2). The binding affinity of spike protein with ACE2 represents important determinant of coronavirus host range ^14,15,16^. Spike proteins are trimers containing two functional subunits designated as S1 and S2. S1 is responsible for binding to the receptor and S2 is responsible for fusion of cellular membrane ^17^. The interacting residues between pathogen and host proteins can be identified using molecular docking technique. It may provide the scope to characterize the more evolutionary constrained regions as target in the pathogen to avoid rapid drug and vaccine escape mutants. Various mutation sites for spike proteins from SARS-CoV-2 isolates have been mapped on to protein three-dimensional structure ^18^. It is reported that the spike protein of SARS-CoV-2 is in a highly stable state and binds to the ACE2 with the higher affinity ^19^.

The huge pool of genome sequence data of SARS-CoV-2 provides us ample opportunity to analyze the evolutionary dynamics of the virus. The rapid spread of SARS-CoV-2 raises many questions on the mutational diversity and its impact on evolution of the virus. The present study is designed to focus on the mutational pattern of spike protein sequences, the underlying cause of variation of mutational patterns and their significance to the binding affinity with human host ACE2 protein. Spike protein has been studied as a potential drug target and also as a virus antigen ^20^. Therefore, the present study might be important for the development of therapeutic, and prevention of SARS-CoV-2.

## Materials and Methods

### Sequence retrieval and analysis

A total number of 251430 whole genome sequence assemblies flagged as “complete”, “high coverage”, “low coverage excl” for human host were downloaded as of January 18, 2021 (8:00 GMT) from GISAID (https://www.gisaid.org/). Full length Spike gene sequences were retrieved through BLAST search by aligning with a reference spike gene sequence (GenBank accession number: NC_045512.2). Those sequences containing unrecognized start codon, stop codon, internal stop codons, untranslatable codons and unrecognized character (other than a,t,g,c) have been discarded from the final dataset. The final set comprises 209148 spike gene sequences.

Correspondence analysis was performed to assess the variations in amino acid usage of spike protein dataset ^21^. CoA reveals major trends of variation in the dataset by arranging them along continuous axes where consecutive axis have been arranged to have diminishing effect gradually ^22^. We used CoA available in CodonW for the analysis of amino acid usage of spike gene sequences. Hydrophobicity of each spike gene sequence is calculated using the method Kyte-Doolittle (1982) available in CodonW ^23^.

### Protein homology modeling and docking

We have followed the method proposed by Huang et al. (2010) ^24^ for clustering of all the spike protein sequences on the basis of sequence identity. We found 7883 clusters of spike proteins with 100% sequence identity. Therefore, we took one spike protein sequence from each cluster for structural analysis. Out of 7883 protein sequences, 3676 spike proteins belong to SPMHR (Spike proteins with More Hydrophobic Residues) and 4207 belong to SPLHR (Spike proteins with Less Hydrophobic Residues) group.

Three dimensional structural models were generated for spike protein sequences through homology modeling. Spike protein structure available in Protein Data Bank (PDB) (PDB ID: 6VYB) was used as template for homology modelling with more than 99% sequence identity and 94% query coverage. The structure of angiotensin converting enzyme 2 (ACE2) was also generated through homology modelling using the protein sequence available in UniProt (UniProt ID: Q9BYF1) and then a template was used from PDB (PDB ID: 6M18) with 100% sequence identity and 99% query coverage. 3Drefine web-server was used for the refinement of the protein structural models generated through homology modeling ^25^. Molecular interaction of viral spike protein with the human ACE2 receptor was performed using Z-dock software ^26^. Then, the resulting docking data were processed and analyzed considering binding energies and main interacting residues in each complex by using the tools of PRODIGY software ^27^.

## Results

### 1. Correspondence analysis on amino acid usage of spike proteins

We analysed the variations in the amino acid usage patterns of spike gene sequences through Correspondence Analysis. Correspondence Analyses are used to simplify rectangular matrices in which (for our purpose) the columns represent amino acid usages value and the rows represent individual genes. It creates a series of orthogonal axes to identify trends that explain the data variation, with each subsequent axis explaining a decreasing amount of the variation. Figure 1 shows the positions of the genes generated through Correspondence Analysis on amino acid usage along the first and second major axes. The first and second major axis accounted for 29.63% and 12.81% of the total variation in amino acid usage respectively. Since there exists a single major explanatory axis (i.e., horizontal axis with 29.63% variation) we, therefore, carried out remaining analysis in this study on the basis of the distribution of spike protein genes along the horizontal axis of Correspondence Analysis. It was observed that the position of the genes along the horizontal axis (Figure 1) were significantly correlated with the hydrophobicity of the encoded proteins (r = 0.45, p < 0.01). We have also observed that average hydrophobicity of the spike proteins distributed in the negative side of the horizontal axis is significantly lower (P<0.0001) than the average hydrophobicity of the spike proteins distributed in the positive side of the horizontal axis. For lucidity, henceforth, spike proteins distributed in the positive side of the horizontal axis will be referred to as Spike Proteins with More Hydrophobic Residues (SPMHR) and spike proteins distributed in the negative side of the horizontal axis will be referred to as Spike Proteins with Less Hydrophobic Residues (SPLHR).

**Figure 1:**
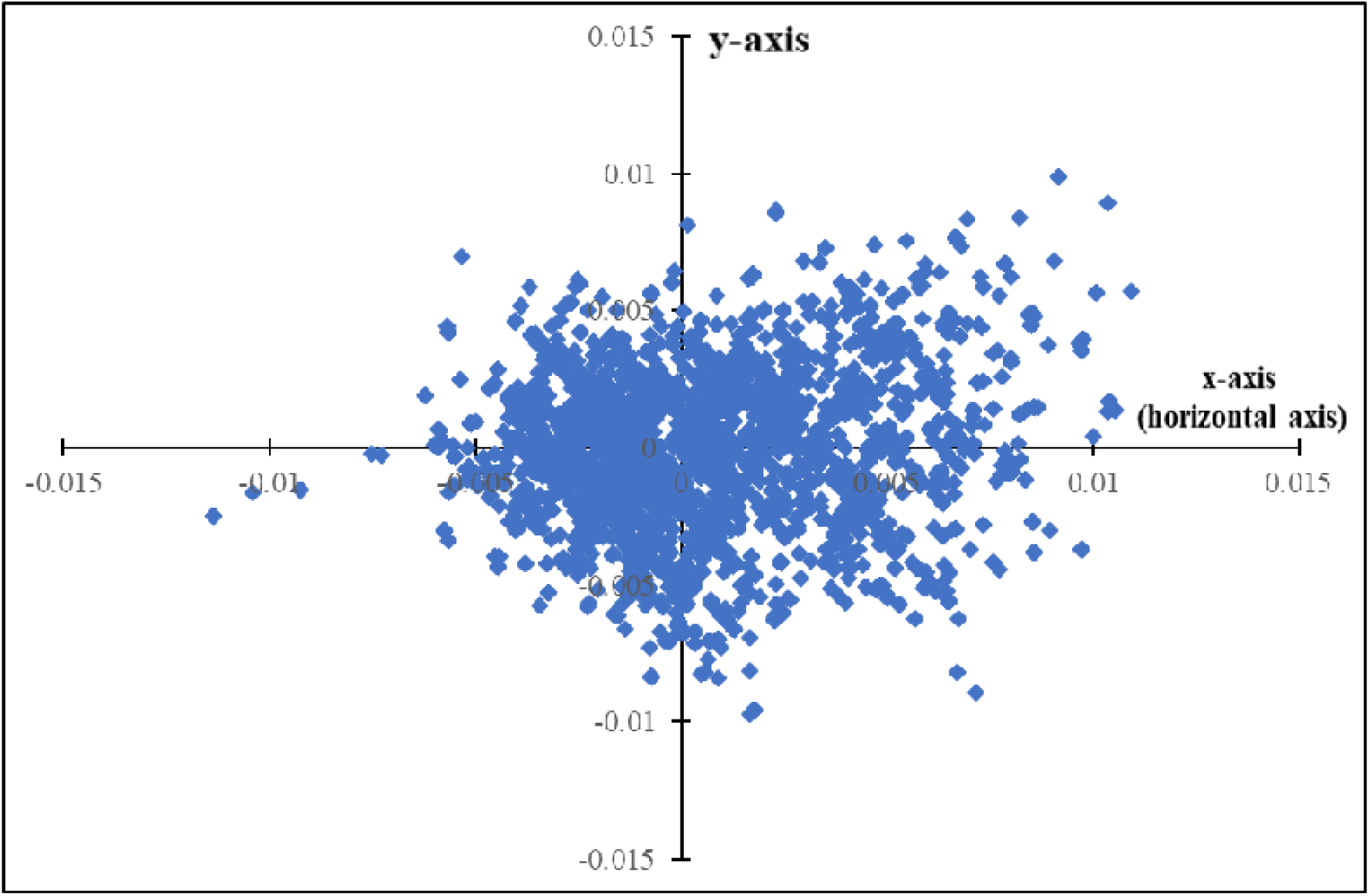
Distribution of Spike(S) genes along the two major axes of Correspondence analysis (COA) based on amino acid usage (AAU) data. Blue coloured square boxes represent Spike(S) gene sequences.

### 2. Distribution of date of sample collection according to differential pattern of spike protein sequences

We have compared the sequence dataset for SPMHR and SPLHR and checked the distribution of date of sequence collection for every spike protein in both the groups. We observed a skewed distribution of spike genes between SPMHR and SPLHR groups with respect to the date of collection of sequences (Figure 2). For the first couple of months (December, 2019 – September, 2020) the percentage of sequences clustered in SPLHR group is higher than the percentage of sequences clustered in SPMHR group. However, from October, 2020 the percentage of sequences clustered in SPMHR group became higher than the percentage of sequences clustered in the SPLHR group. Interestingly, as we move from December, 2019, the percentage of spike protein sequences clustered in SPLHR gradually decreases with an increase in the percentage of spike protein sequences clustered in SPMHR group. To our surprise, we observed that almost 100% of spike protein sequences in December, 2019 followed SPLHR pattern whereas, almost 50% of spike protein sequences followed SPMHR pattern from October, 2020.

**Figure 2:**
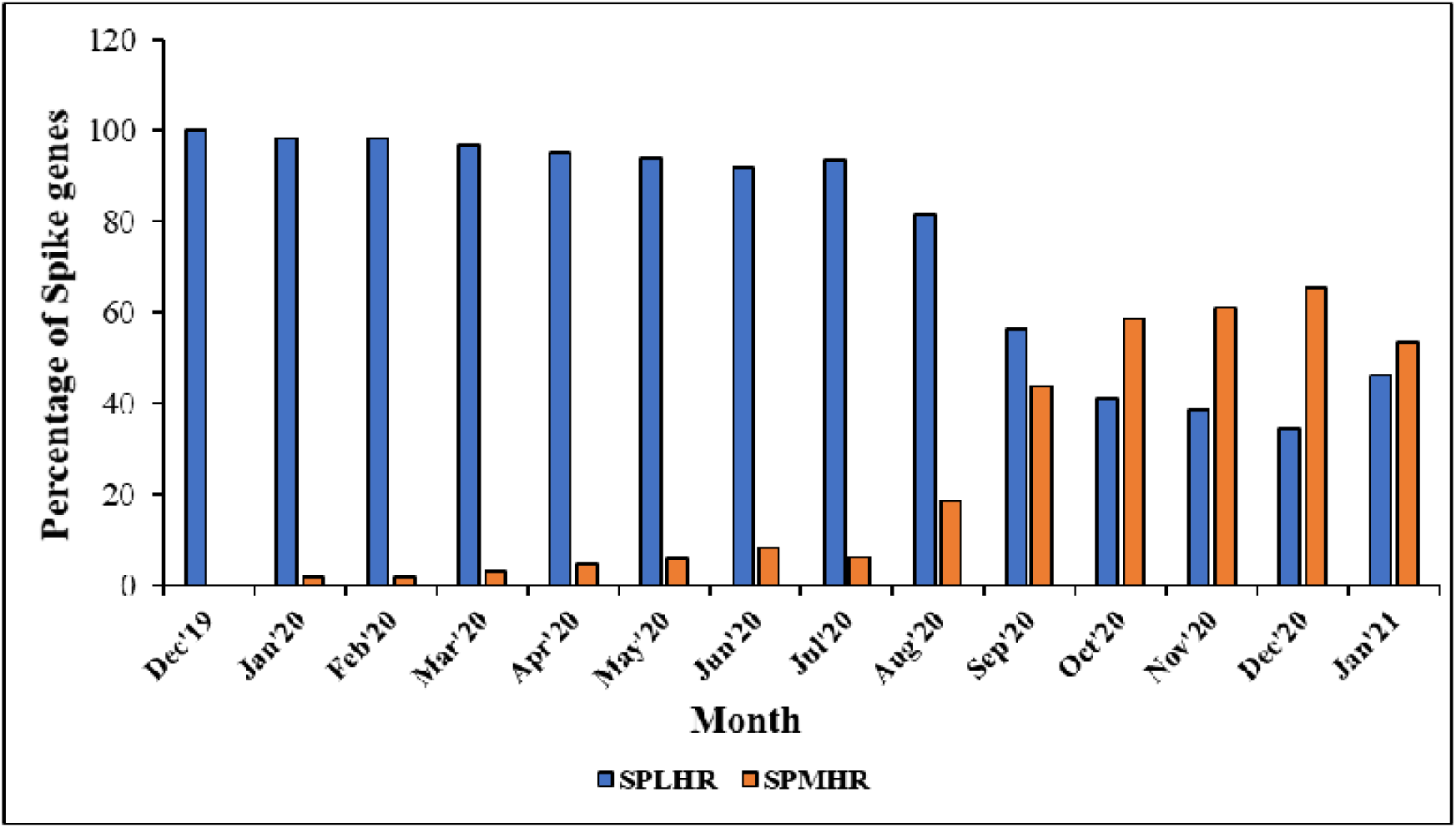
Distribution of spike genes between SPMHR and SPLHR groups with respect to the date of collection of sequences.

### 3. Interaction profile between spike protein and ACE2

Our comparison of amino acid usage underlines the differential pattern of evolution of Spike protein where the hydrophobicity of the encoded protein is the major cause of variation of amino acid usage pattern among the spike proteins. Since the receptor for SARS-CoV-2 has been identified as ACE2 it was very important to analyze how the differential pattern of amino acid usages of Spike proteins of SARS-CoV-2 responded to binding to the human ACE2 receptor. Three dimensional structures of Spike protein sequences from the SPMHR cluster and SPLHR cluster were constructed through homology modeling using the crystal structure of Spike protein available in PDB (PDB ID: 6VYB) as template. The 3D structure of ACE2 has been generated computationally using the protein sequence available in UniProt (UniProt ID: Q9BYF1) and then a template was used from PDB (PDB ID: 6M18). Docking study was performed with ACE2 separately with all the spike proteins taken from the two groups (Figure 3(A)) and the average binding energy was calculated separately for SPMHR and SPLHR groups. We observed that the average binding energy for the spike-ACE2 complex taken from the SPMHR group is significantly lower than the average binding energy for the same complex taken from the SPLHR (P<0.0001). A lower average binding energy for the spike-ACE2 complex taken from the SPMHR indicates its higher stability compared to the SPLHR group of complexes.

**Figure 3(A):**
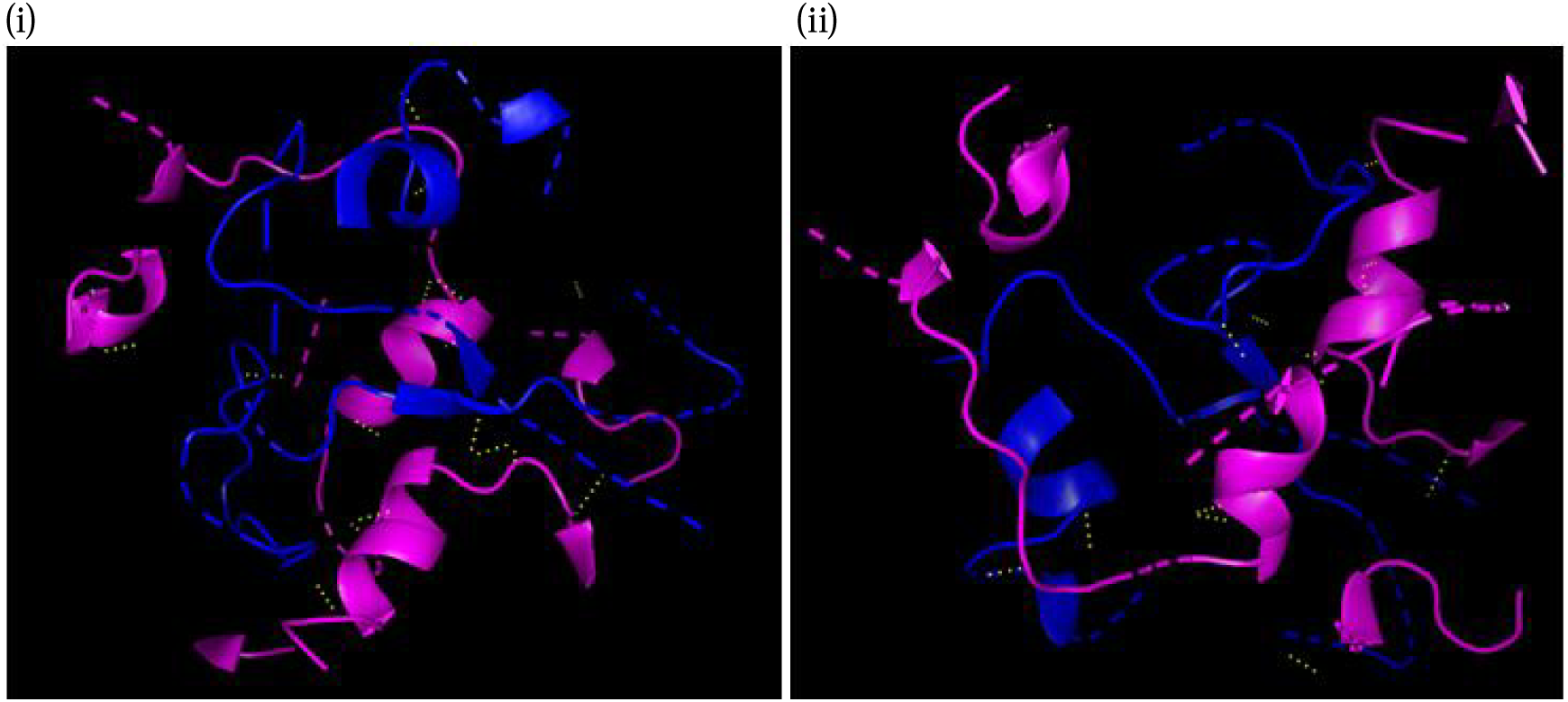
Arrangement of hydrogen bonds (yellow) in Spike (Blue)-ACE2 (Pink) complex. (i). Spike protein taken from the SPMHR indicates ACE2 receptor can interact with viral spike protein more effectively compared to (ii). Spike-ACE2 complex where spike protein was taken from the SPLHR group.

**Figure 3(B):**
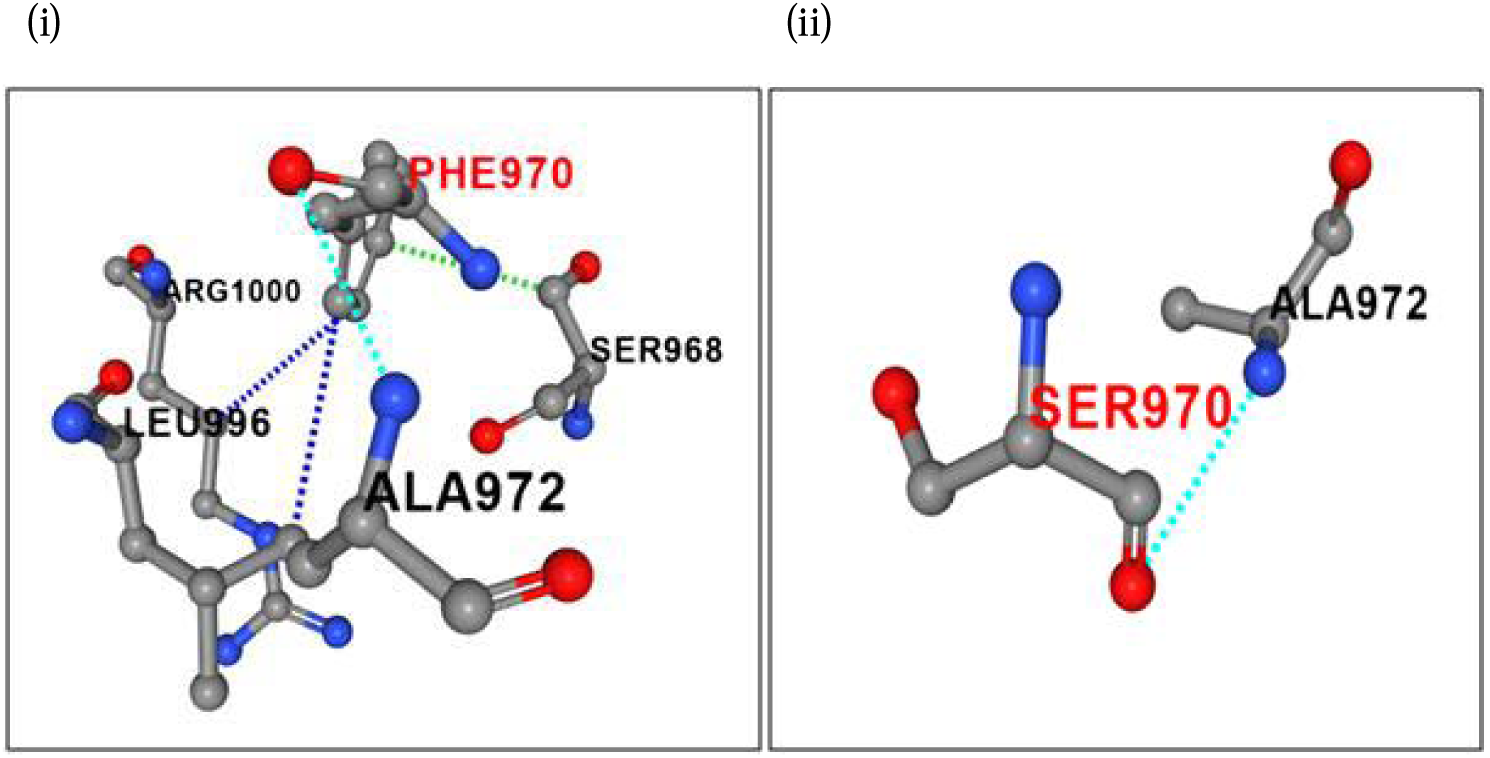
Comparison of interaction profile of an identified mutation F970S in Spike-protein indicating hydrophobic to hydrophilic amino acid substitution. (i) Wild type residue F970 having one polar interaction (sky), one Van der Waals interaction (green) and two hydrophobic (blue) interaction. (ii) Mutant type residue 970S having one polar interaction (sky).

## Discussion

We performed a comprehensive analysis of amino acid usage of more than 0.2 million spike proteins to understand the evolutionary dynamics of the emerging SARS-CoV-2 pandemic. Earlier studies have identified several mutations in the Spike protein ^10,28,29^; however, the present study has categorically analyzed the underlying cause of different kinds of mutations that has shaped the evolution of Spike protein during the ongoing pandemic. In the present study, variation of hydrophobicity of Spike protein was observed to be the most important factor influencing the amino acid changes in spike protein. It is argued that the majority of viral mutations is harmless, however, some of these mutations may change infectivity, survival capability, pathologic property, or immunogenicity and antigenicity of the virus ^30,31^. Previous reports pointed out several important mutations and in most of these cases directional mutational pressure changed towards higher hydrophobic/lower hydrophilic amino acid ^9,32,33^. Recent studies suggest that the D614G variant is close to reaching fixation around the world ^12^. The present work reinforces this observation that changes between Aspartic acid to Glycine contributes towards the difference in amino acid usage of Spike protein associated with a change in hydrophobicity. The mutation D614G on the spike protein may also increase binding affinity between the spike protein and host ACE2 receptor, thus enhancing virus loads which lead to increased infectivity ^34^. In addition, recent reports also provide evidence of this variant in viral spread associated with higher viral load^35,36^.

Our results also show that compared to spike proteins collected during the earlier months the recent sequences preferred to have more hydrophobic residues, as is evident from the higher number of sequences followed SPMHR pattern (Figure 2). The functional significance of lower binding energy for the spike-ACE2 complex for SPMHR group indicates that the ACE2 receptor can interact with viral spike protein more effectively compared to the spike-ACE2 complex from the SPLHR group. To determine the potential role of hydrophobic residues towards the binding affinity of Spike-ACE2 complex, we identified how many residues have changed between two proteins taken for our molecular docking study. We observed six mutations (V405D, A414Q, G614D, V860Q, L861K, F970S (Table:1) that revealed hydrophobic to hydrophilic amino acid substitution. Figure 3(B) shows the change in the bonding pattern when phenylalanine was substituted with serine. The Gibbs free energy for unfolding was calculated to evaluate the effects of each of these mutations individually on protein complex stability^37^. The positive change in free energy associated with five of these mutations indicated destabilizing mutations. Only A414Q was linked with a negative free energy change, however, the free energy value was negligible and least among all the mutations considered (Table 1). Therefore, accumulation of hydrophilic amino acid by substituting hydrophobic amino acid has a destabilizing effect on the Spike-ACE2 complex. Selection of more hydrophobic amino acids in Spike protein has functional significance on more effective binding with ACE2 receptor ^34^.

**Table 1:**
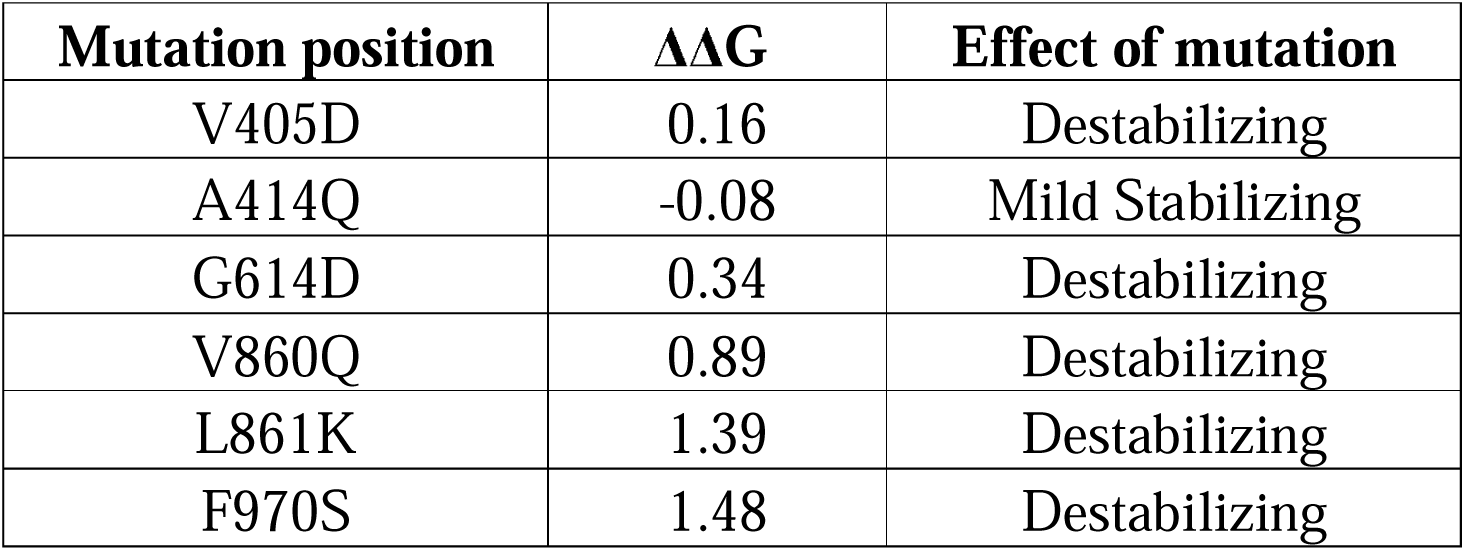
Unfolding Gibbs free energies of mutations calculated using PremPS server depicting change in protein complex stability upon mutation.

| **Mutation position** | **$\Delta\Delta G$** | **Effect of mutation** |
| --- | --- | --- |
| V405D | 0.16 | Destabilizing |
| A414Q | -0.08 | Mild Stabilizing |
| G614D | 0.34 | Destabilizing |
| V860Q | 0.89 | Destabilizing |
| L861K | 1.39 | Destabilizing |
| F970S | 1.48 | Destabilizing |

## Conclusion

The present study shed light on the protein hydrophobicity as the mutational pressure for the evolution of spike protein of SARS-CoV-2. Here, in the present manuscript, we have provided a comprehensive analysis of molecular evolutionary data to understand the virus infection potential which might be important for public health measures and prevent future epidemics like SARS-CoV-2. However, it would be judicious to consider the possibility that mutational variants might modulate the virulence and thereby might have impact on the pathogenicity of the disease. The classification of spike proteins according to the variation of hydrophobicity and thereby modulating the receptor binding affinity provides crucial information for designing treatment and, eventually, vaccines. The findings of the present study could help for the design of potential vaccine candidates/small molecular inhibitor against COVID19.

## Acknowledgement

Manisha Ghosh is supported by Senior Research Fellowship by Indian Council of Medical Research (ICMR).

